# Older Adults Show Altered Default Mode and Executive Control Network Connectivity during Fairness Decisions

**DOI:** 10.1101/2025.08.13.670194

**Authors:** Dominic S. Fareri, Yi Yang, Rita M. Ludwig, James B. Wyngaarden, Derrick Dwamena, Katherine Hackett, Tania Giovannetti, David V. Smith

## Abstract

Fairness-related decisions depend on both offer value and partner identity, yet how aging reshapes the network-level connectivity supporting these decisions across social contexts remains unknown. Younger (18–35) and older (65–80) adults underwent fMRI while responding to offers in an ultimatum game from ostensible partners similar or dissimilar in age. We modeled trial-by-trial choices by offer size, age group, and partner similarity, and examined whole-brain activation and task-dependent connectivity of the default mode (DMN) and executive control (ECN) networks. Participants accepted fairer offers more often, but behavioral sensitivity did not differ reliably by age group or partner similarity. Whole-brain activation analyses revealed no significant age-by-partner-similarity effects. Connectivity analyses, however, revealed opposing age-related patterns. Younger adults showed stronger fairness-modulated DMN–anterior cingulate connectivity for similar than for dissimilar partners, whereas older adults showed the reverse. Age also moderated the relationship between individual sensitivity to fairness norm violations and ECN–medial prefrontal connectivity: this association was negative in younger adults and positive in older adults. These findings suggest that decisions regarding fairness in younger and older adults may be supported by different patterns of large-scale network connectivity.

## Introduction

Financial insecurity and fraud are increasing among older Americans (Collins & Casey, 2017), presenting a significant public health problem. Among the most common types of scams targeting older adults are those that attempt to exploit social identity and relationships –e.g., a scammer may pretend to be a grandchild calling in distress. Such impersonation scams cause a loss of approximately 8 billion per year in older victims (Johnston, 2025). Insights from neuroscientific research of aging-related brain changes during information processing, learning, and decision-making may help develop better protections for older people against scams and fraud. The present research adds to a growing body of literature on financial decision-making in older adults by using a multi-trial ultimatum game with varying social context conditions related to age group similarity between participants and their game partners. The goal of this work is to illuminate associated neural structures and networks implicated during fairness-related choice responses within this game and identify distinctions in the effects of partner similarity and individual sensitivity to norm violations between age groups.

Given the real-world influence of social context on financial decisions in older adults, we implemented an ultimatum game with multiple social conditions (e.g., game partner identity) to study such decision-making in the lab. The ultimatum game provides a powerful tool for studying fairness-related decision making, as it is a take-it-or-leave-it class of economic games wherein a proposer divides an endowment with a responder, who then either chooses to accept or reject the offer. If the responder rejects the offer, both parties walk away with nothing (Güth et al., 1982). These games are thought to engage multiple processes including social cognition, reward valuation, and learning by provoking a motivational conflict between economic self-interest and higher-order social goals of maintaining norms of fairness and reciprocity (Andreoni & Bernheim, 2009). Indeed, despite the economically rational choice being to always accept non-zero offers to increase personal net gain, responders frequently make the pyrrhic choice to reject unfair offers (Güth & Kocher, 2014).

By manipulating age similarity between a player and their game partner, we investigated how social contextual factors influenced the player’s choice to accept or reject an offer and related brain activations, and whether these behavioral and neural patterns differed between younger and older adults. At the behavioral level, research on age differences in the ultimatum game has yielded conflicting results; older adults have been reported to both accept more unfair offers than younger adults (Fernandes et al., 2019; Girardi et al., 2018) and, conversely, to reject them more frequently (Harlé & Sanfey, 2012). At the neural level, neuroimaging research has focused on identifying distinct regional brain activations and has shown age differences in the dorsolateral prefrontal cortex, anterior insula (e.g., Harlé & Sanfey, 2012; Gao et al., 2018), and ventromedial prefrontal cortex (e.g., Gu et al., 2015).

The focus on activation of specific regions overlooks the distributed nature of neural processing and the role of large-scale brain networks (Feng et al., 2021; S. M. Smith et al., 2009) in social decision-making, which because of its complexity depends on coordinated interactions across multiple neural systems (Ruff & Fehr, 2014; Stanley & Bharat, 2020). Because aging alters communication within neural networks supporting goal-directed behavior (Andrews-Hanna et al., 2007; Madden et al., 2010; Schlesinger et al., 2017), it is crucial to begin characterizing how age-related changes in network connectivity may be associated with behavior during fairness-centered social interactions.

Here, we focus on two networks that have been repeatedly implicated in aging, social cognition, and decision making: the default mode network (DMN) and the executive control network (ECN). The DMN, which includes regions such as the medial prefrontal cortex, posterior cingulate cortex, and temporoparietal junction, has been linked to self-referential thought, social cognition, and the subjective valuation of social outcomes (Andrews-Hanna et al., 2014; Buckner & DiNicola, 2019; D. Smith et al., 2026). Importantly, DMN connectivity is sensitive to social context during trust-based interactions, showing enhanced coupling with reward-related and attentional-control regions when individuals experience reciprocity from close others versus strangers (Fareri et al., 2020). The ECN, which encompasses lateral prefrontal cortex, anterior cingulate cortex, and anterior insula (S. M. Smith et al., 2009), supports goal-directed behavior, cognitive control, and the regulation of prepotent responses, all functions that are critical when navigating conflicts between self-interest and social norms during economic exchanges (Madden et al., 2010; Waltz et al., 2013). Recent work from our group found that older adults show altered ECN connectivity with insular cortex during trust game outcomes, suggesting that age-related changes in ECN function may influence how social information is processed in economic contexts (Fareri et al., 2022). By examining task-dependent connectivity of both networks as a function of age and social context, we aim to clarify how fairness decisions are supported across the lifespan and whether younger and older adults exhibit different connectivity patterns during fairness-based social interactions.

Importantly, not all people will respond to violations of fairness norms in the same way. For example, some individuals tolerate substantial deviations from equal splits, while others reject even modestly unfair proposals (Strobel et al., 2011; Thielmann et al., 2016). This variability reflects stable individual differences in sensitivity to fairness norm violations, which have been linked to distinct patterns of activation in regions supporting social evaluation and cognitive control, including the anterior insula, dorsolateral prefrontal cortex (dlPFC), and anterior cingulate cortex (ACC) (Gabay et al., 2014; Sanfey et al., 2003). Such individual differences may interact with both age and social context. Older adults have been shown to weigh outcome equity and proposer intentions differently than younger adults (Cho et al., 2020), and sensitivity to social norms may be modulated by whether one is interacting with an in-group or out-group partner (Baumgartner et al., 2012). Thus, by examining how individual-level sensitivity to norm violations relates to neural connectivity during the ultimatum game, we can begin to move beyond group-level effects to understand how stable behavioral dispositions toward fairness are instantiated in the brain and whether these brain-behavior relationships differ across age groups and social contexts.

The present study sought to clarify and extend previous mixed extant findings on the neural processes supporting social decision-making in older adults during the ultimatum game. We included a social context manipulation that went beyond a human versus computer partner paradigm (e.g., Batten et al., 2024; Harlé & Sanfey, 2012) by including two purported human partners to manipulate partner similarity based on age, which has been shown to relate to judgments of interpersonal trust and offer rejection rates (Alarcon et al., 2016; Bailey et al., 2013). We hypothesized that when interacting with human partners, younger adults would show an increased tendency to accept fair offers when interacting with age-similar vs. age-dissimilar partners. We also expected that older adults’ tendency to accept fair offers would not differ according to partner similarity. At the neural level, we investigated whether neural responses and connectivity to unfair offers differed across age groups and partner types. Based on prior empirical and meta-analytic work (e.g., Fehr & Camerer, 2007; Gabay et al., 2014), we hypothesized that neural activation in the anterior insula and dlPFC and ACC would be diminished in older adults during receipt of unfair offers from age-similar vs. age-dissimilar partners. We also hypothesized that age-dependent changes in neural responses to offers will be tied to differential changes in connectivity of both the DMN and ECN. Furthermore, we explored whether patterns of brain activity were related to an individual’s behavioral sensitivity to fairness norm violations.

## Methods

### Participants and Procedure

We recruited 50 participants into two groups of interest, younger adults aged 18-35 (N = 26) and older adults aged 65-80 (N = 24). Our sample size was planned *a priori* to collect at least 20 usable participants in each group for a total of at least 40 participants; this planned sample size was limited by available funding for data collection. Young adult participants were recruited primarily through the Temple University Psychology Department participant pool, while older adult participants were recruited locally in Philadelphia, PA via posted flyers and newspaper advertisements, and reaching out to community and senior centers. All participants were screened before data collection to rule out major psychiatric or neurologic illness as well as MRI contraindications. Older adults were screened to rule out dementia using the Telephone Interview for Cognitive Status (Brandt et al., 1988). Participants were further excluded if their average framewise displacement in their brain images exceeded the upper bound defined by the boxplot method (1.5 × Interquartile Range above the third quartile). This sampling procedure resulted in a final sample of 47 participants consisting of 25 young adults (mean age = 23.40 years, SD = 4.01; 16 female, 9 male) and 22 older adults (mean age = 69.32 years, SD = 4.36; 11 female, 11 male). Given our final sample (n = 25 younger adults; n = 22 older adults), we conducted a sensitivity analysis in G*Power 3.1 for a two-tailed independent-samples t test (alpha = .05, power = .80). This analysis indicated that our design had 80% power to detect age-group differences of approximately d = 0.84, which corresponds to approximately R^2^ = .15 and f^2^ = 0.18 for an equivalent regression model with a single group indicator. Smaller true effects could still be observed in our data, but they would be detected with less than 80% power.

Participants recruited through the participant pool were compensated with course credit, while those from the community were compensated $25 per hour of participation with Amazon and/or Visa gift cards; to increase the ecological validity of the task, we additionally provided bonus payments based on one randomly chosen trial from the experimental session. This study was approved by the Temple University Institutional Review Board and all participants gave informed consent.

After assessing eligibility based on initial screening procedures, participants were invited to participate in our study. We collected various measures as part of a larger neuroimaging experiment (see Smith et al., 2024 for more details), but here we will report only the pertinent details for the ultimatum game. Participants completed two appointments. During the first appointment, which lasted approximately 90 minutes, they underwent a mock MRI scan to acclimate them to the scanner and help control for motion. They then completed a brief neuropsychological test battery, with older adults additionally completing a self-report measure of everyday functioning. At the beginning of the second appointment, participants played a practice version of the ultimatum game to ensure understanding of the task and to reduce the likelihood of missed trials during the scan. The remaining duration of this appointment was spent playing the ultimatum game task and other economic games in the scanner and completing post-scan tasks (not reported here), after which participants were debriefed, compensated, and dismissed.

### Experimental Design

Participants played as responders in a multi-trial version of an ultimatum game in which we manipulated both the fairness of proposed offers and the social context in which they were presented. Participants received 144 total offers (18 blocks with 8 trials each) from three partners with whom they were told they would be splitting a $20 endowment: a computer (which served as our non-social control), a gender- and race-matched younger adult, and a gender-and race-matched older adult. Whether the human partner was considered a ‘similar’ or ‘dissimilar’ condition was therefore dependent upon the age of the participant. On each trial (see Figure 1), participants saw a screen with the offer value and the partner proposing that split as cued by a cartoon image of a younger adult, older adult, or computer (non-social control condition beyond the scope of the current investigation and excluded from analyses). Fairness was operationalized by the equity of the offer size proposed to the participant: offers were either mostly fair (i.e., between 35% – 50% of the endowment value) or unfair (between 5% – 20%). Following an interval of one second, participants had 2.5 seconds to either reject or accept the offer.

**Figure 1.**
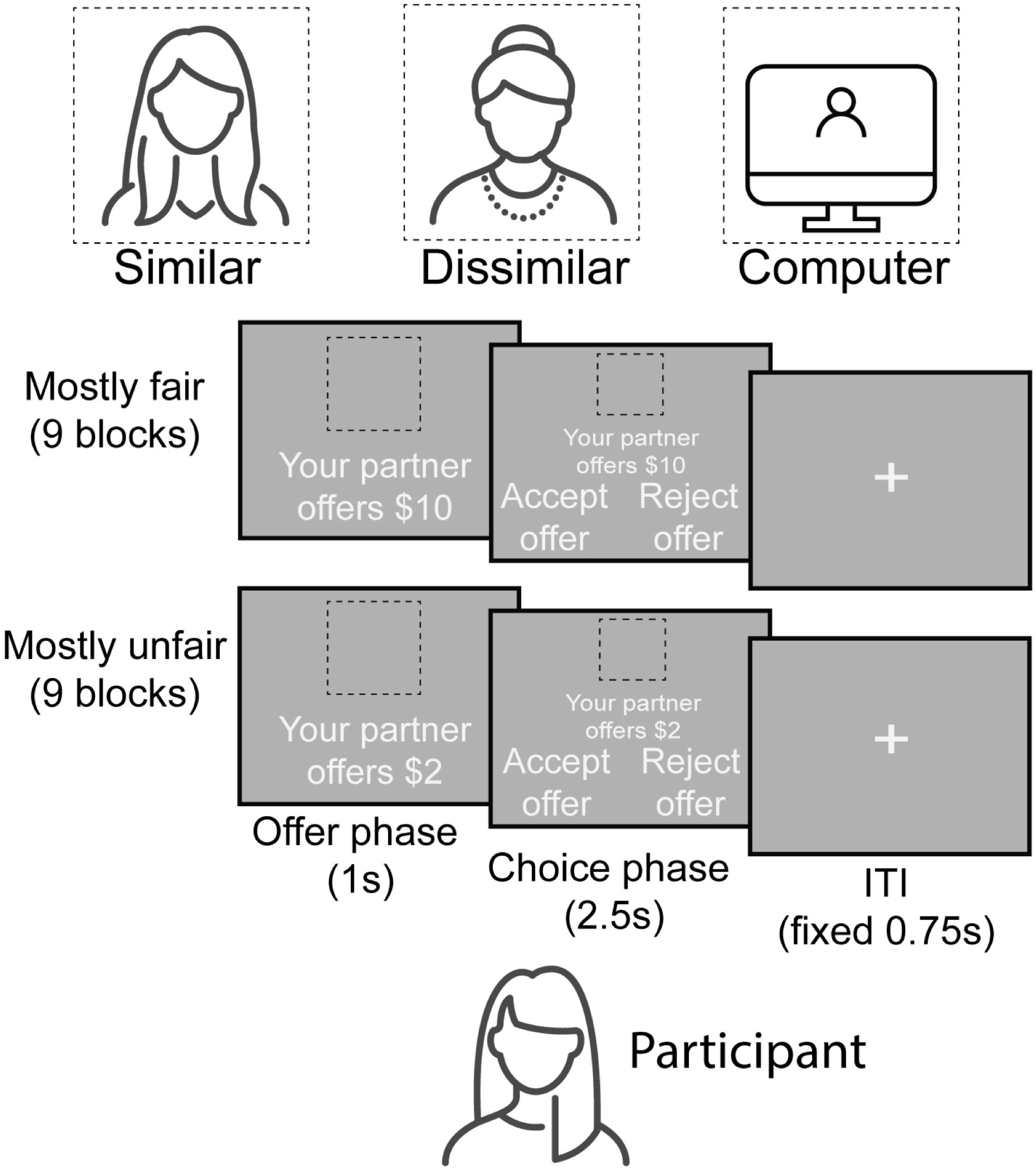
Ultimatum game task. Participants received 144 total offers (18 blocks with 8 trials each) from three partners with whom they were told they would be splitting a $20 endowment: a computer (non-social control beyond the scope of the current investigation and excluded from analyses), a gender- and race-matched younger adult, and a gender- and race-matched older adult. Whether the human partner was considered a ‘similar’ or ‘dissimilar’ condition was therefore dependent upon the age of the participant. Each offer was either mostly fair (between 35% – 50% of the endowment value) or unfair (between 5% – 20%). Condition types of offer size and partner similarity were fully crossed, such that participants saw all offer sizes from all three partner types, though the order of the pairing of these conditions was randomized.

### Neuroimaging Data Collection and Preprocessing

Neuroimaging data were collected as part of a larger pilot study involving age-related differences in social and economic decision-making. Functional data were collected with a spatial resolution of 2.97×2.97×2.80 mm with a 2.02 s repetition time. All imaging data were first converted to BIDS format using HueDiConv (Halchenko et al., 2024) before further processing. Preprocessing of neuroimaging data was performed using fMRIPrep 20.2.3 (RRID:SCR_016216; (Esteban et al., 2019), which is based on Nipype 1.6.1 (RRID:SCR_002502; (Gorgolewski et al., 2011). Details of fMRI image acquisition, fMRIprep processing, and the full dataset have been reported in our protocol paper and are openly available on OpenNeuro (D. V. Smith et al., 2024).

### Behavioral Analyses

To test our behavioral hypothesis, we employed a hierarchical model comparison approach using a series of multilevel logistic regression analyses on the trial-by-trial choice data (accept or reject) (Table 1). This method was selected to account for the nested data structure and to systematically identify the most parsimonious model.

**Table 1.**
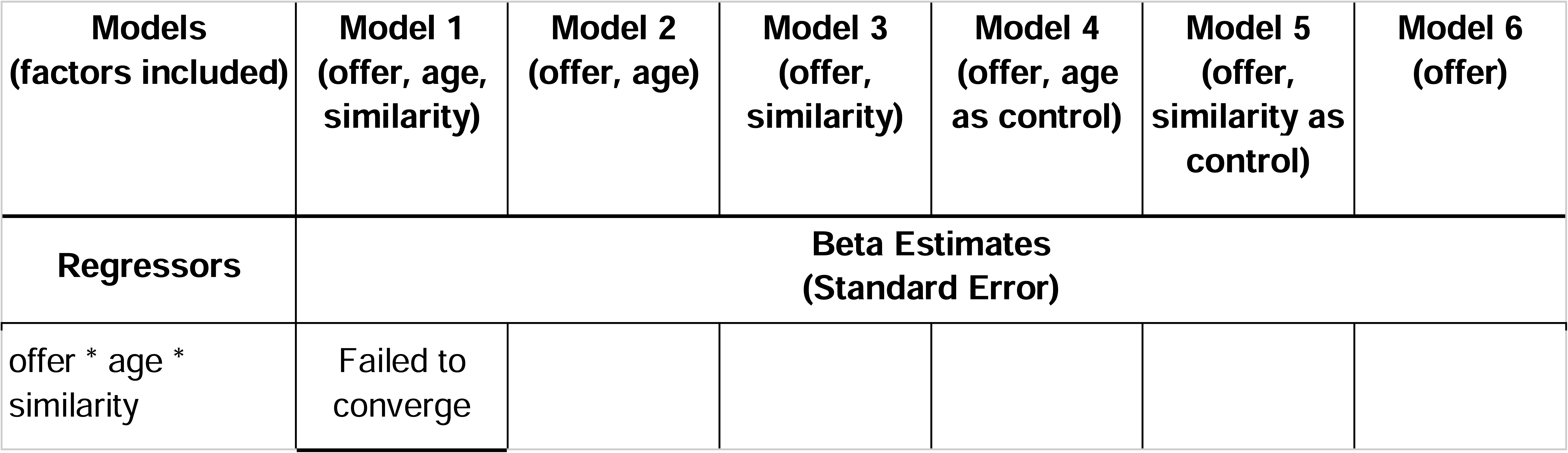
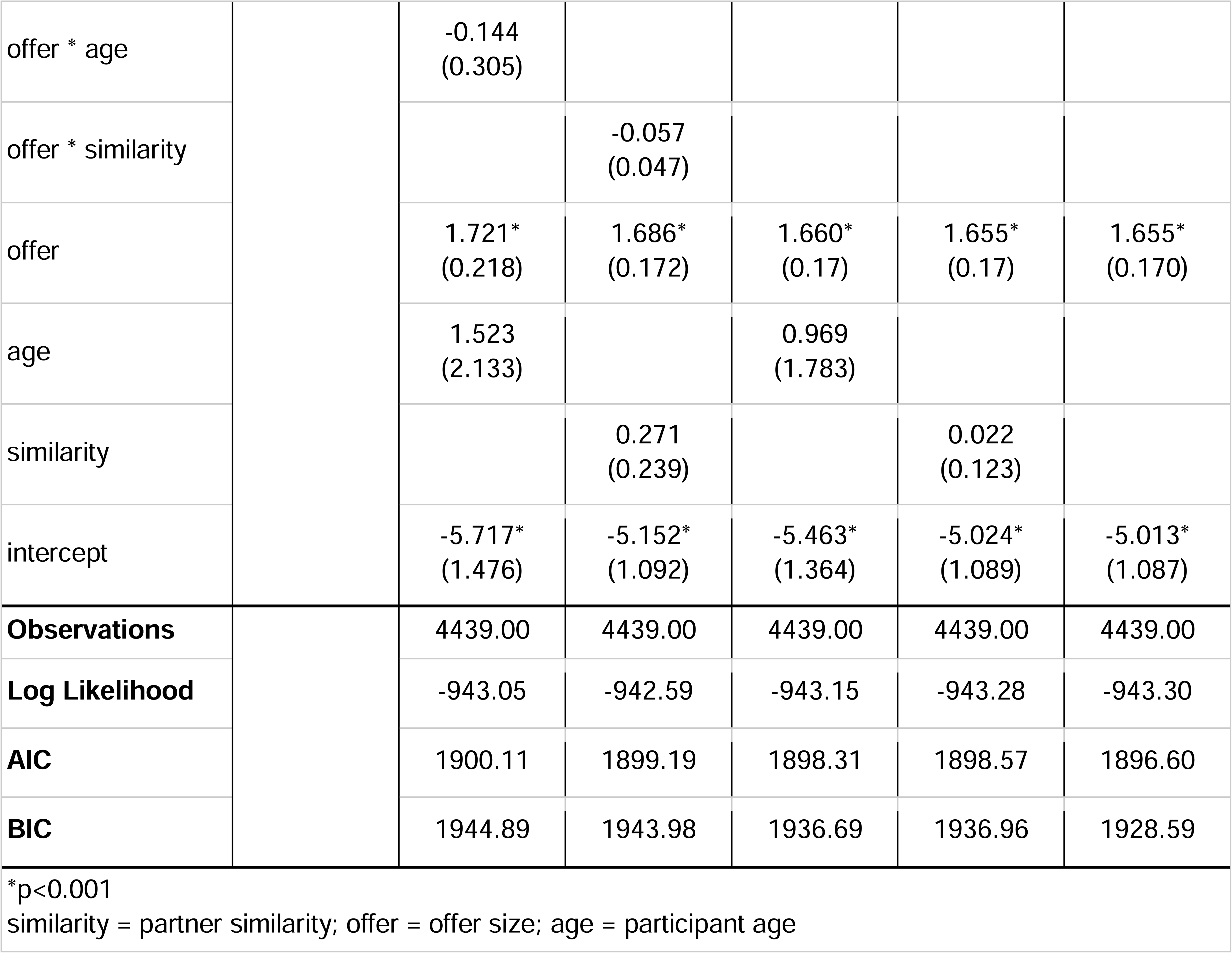
Models and Estimates (fixed effects, *p<0.001)

Our analyses began with a full model predicting the likelihood of offer acceptance. This initial model included fixed effects for offer size, partner similarity, participant age group (younger vs. older), all two-way interactions, and the critical three-way interaction among these variables (Model 1 in Table 1). If the three-way interaction was not significant, we proceeded by testing a more simplified model that included only the two-way interactions and the main effects (Models 2 & 3). If the two-way interactions were also non-significant, we would then examine a final model containing only the main effects of offer size (Model 4), partner similarity (Model 5), and age group (Model 6) after controlling for offer size. For all models, we included random intercepts and random slopes of offer size for each participant to account for individual baseline differences in decision-making. All behavioral analyses were conducted in R version 4.3.2 with the lme4 package (Bates et al., 2015).

### Quantifying Behavioral Sensitivity to Fairness Norm Violations

To investigate how brain activity relates to individual differences in behavioral sensitivity to fairness norm violations, we first quantified this behavioral tendency for each participant. We derived an individual-level metric from a series of multilevel logistic regression models. For each partner-similarity condition (e.g., similar-age partner, dissimilar-age partner), we fit a multilevel logistic regression model predicting acceptance from offer size, which indexed fairness because all offers were at or below an equal split. the association between choosing to accept and fairness (i.e., offer size). We included both fixed and random intercept and slope of offer size in the multilevel models. From each model, we extracted an individual-specific slope for every participant, which was calculated as the sum of the fixed-effect slope of the offer size and the participant’s unique random-effect slope for offer size. This value represents that individual’s estimated sensitivity to fairness within that specific social context. To create a single metric capturing the influence of partner similarity, we calculated a difference score by subtracting the sensitivity value derived from the dissimilar partner condition from the sensitivity value derived from the similar partner condition. This final score represents each participant’s partner similarity-related sensitivity to fairness norm violations. This individual sensitivity score and its interaction with participant age were then included as additional covariates in our group-level whole-brain analyses (see Group-Level Whole-Brain Analyses subsection below) to identify neural correlates of this fairness norm related behavioral tendency. All brain-behavior analyses were conducted in R version 4.3.2 with the lme4 package (Bates et al., 2015).

### Neuroimaging Analyses

#### fMRI First-Level Modeling

Functional data were analyzed in FSL using a general linear model (GLM) with local autocorrelation correction (Woolrich et al., 2001). We first focused on task-related activation using a GLM consisting of nine regressors. We used three regressors of partner similarity (human similar, human dissimilar, computer) to model trials (duration = 3.5 s) within each of the partner blocks. We also included parametric modulators for each of these three regressors, where each trial within a block was weighted by the offer size (i.e., dollar value of the offer, which is proportional to fairness given a fixed endowment of $20) on that trial. Given that response time may vary across partner similarity conditions and participants, we included additional regressors modeling the response as a constant term and with a parametric modulator for the response time on each trial. Finally, we included a regressor to account for missed trials. Each of these regressors were convolved with the canonical hemodynamic response function.

Task-dependent changes in connectivity were examined using psychophysiological interaction (PPI) analysis (Friston et al., 1997; D. V. Smith et al., 2016; Smith & Delgado, 2017). Regions exhibiting a significant PPI effect with a seed region or network can be interpreted as showing a context-specific modulation of effective connectivity, though we note that the directionality of such an effect is ambiguous without additional analyses (Friston, 2011). Given our goal of examining DMN and ECN connectivity with other brain regions, we elected to use a network PPI approach that is not limited to single seed regions (Dobryakova & Smith, 2022; Utevsky et al., 2017). We conducted two separate generalized network PPI models using the 10 networks characterized in prior work (S. M. Smith et al., 2004): one with the ECN as the primary network of interest (with the remaining nine networks, including DMN, as covariates) and one with DMN as the primary network of interest (with the remaining 9 networks, including ECN, as covariates). Each network time course was extracted using the spatial regression component of the dual regression approach and then added to the task-related activation model described above. We formed the PPI regressors by multiplying each of the 9 task regressors by the primary network regressor, which produced a total of 28 regressors for each model.

In both activation and connectivity models, we included a common set of confound regressors derived from the output of fMRIprep. Specifically, we included six regressors for head motion parameters (rotations and translations), the first six aCompCor components explaining the most variance, non-steady state volumes, and the framewise displacement across time. We also included a set of discrete cosine basis functions to apply a high-pass filter (128 s cut-off).

#### Group-Level Whole-Brain Analyses

To identify age-related neural responses to offer size, partner similarity, and sensitivity to fairness norm violations, we conducted group-level whole-brain analyses of activation and network connectivity using FLAME (FMRIB’s Local Analysis of Mixed Effects) Stage 1 and Stage 2 (Woolrich et al., 2004). We implemented two types of group-level models focused on the contrast between similar and dissimilar partners (similar partner > dissimilar partner) as the dependent variable. The first model examined the interaction between age and partner similarity and included regressors for age group (younger and older adults) and control covariates of gender, temporal signal to noise ratio (tSNR), mean framewise displacement, and mean response time to control for potential confounding effects of participant demographics and data quality measures. The second model examined the interaction between age, partner similarity, and fairness norm violation sensitivity and included the same covariates as the first model plus two additional regressors for interaction between fairness norm violation sensitivity and participant age group (i.e., norm violation sensitivity to similar minus dissimilar partner x younger adults, norm violation sensitivity to similar minus dissimilar partner x older adults). All z-statistic images were thresholded and corrected for multiple comparisons using an initial cluster-forming threshold of Z > 3.1 followed by a whole-brain corrected cluster-extent threshold of p < 0.05, as determined by Gaussian Random Field Theory (Flandin & Friston, 2019).

## Results

### Relation of Offer Size to Offer Acceptance

For our primary analyses, we first examined if decisions to accept or reject were associated with the fairness of offers, participant age group, and partner similarity to the participant. We found a significant main effect of offer size (Figure 2): participants were more likely to accept an offer as offers became more equitable (beta = 1.65, p < 0.001). Neither age group nor partner similarity moderated this positive association between offer size and offer acceptance (offer size * age group * partner similarity, model did not converge; offer size * age group, beta = -.14, p = .637; offer size * partner similarity, beta = -.06, p = .227. Table 1).

**Figure 2.**
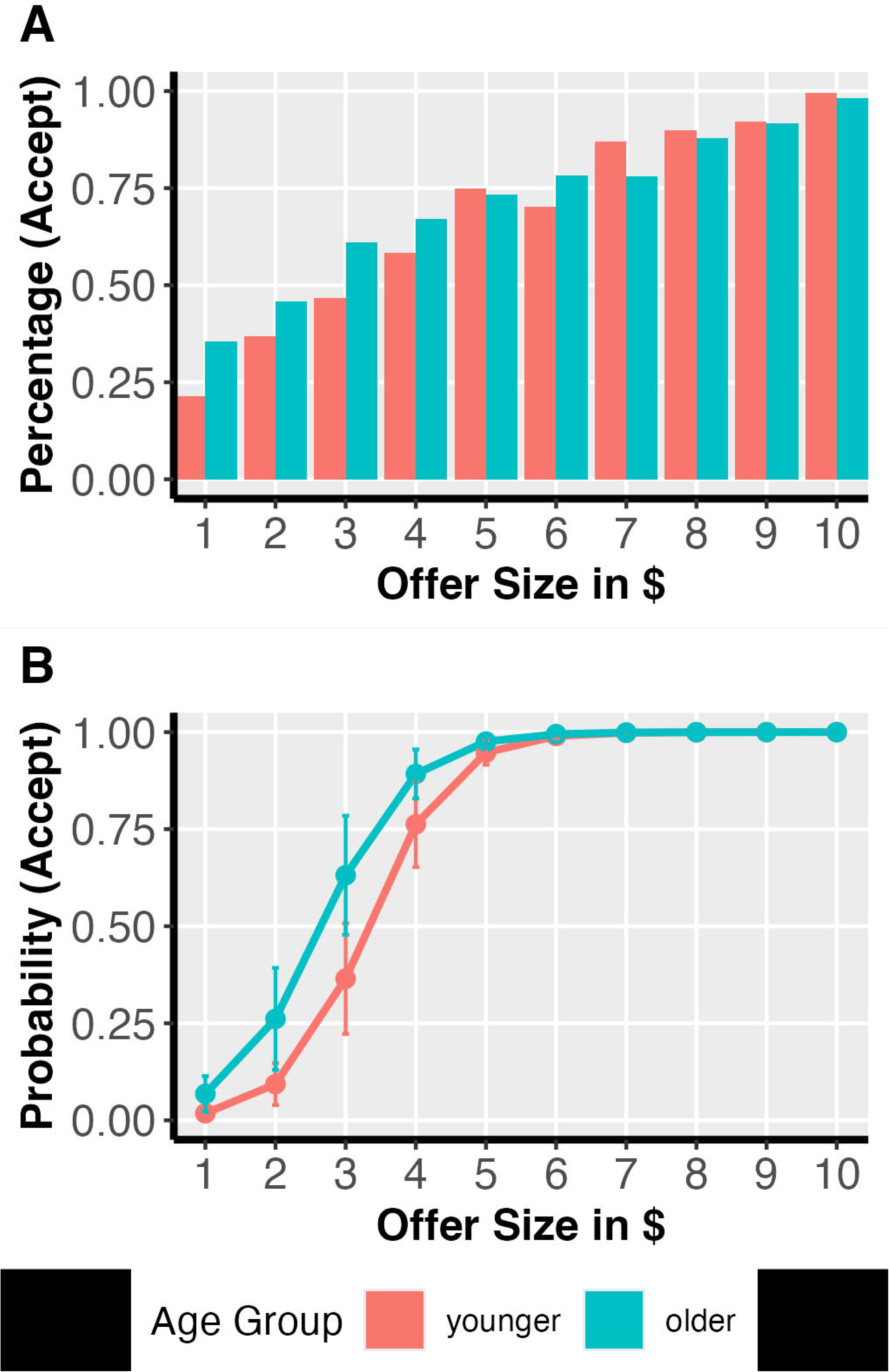
Positive relationship between offer fairness and acceptance probability. The figure displays the probability of participants accepting across offer size. The x-axis represents the size of the offer, indicating its level of fairness, while the y-axis shows the percentage of acceptance (Panel A) and mean probability of acceptance (Panel B). Error bars represent the standard error of the mean for each offer size. A multilevel logistic regression analysis revealed a significant main effect of offer size (beta = 1.65, *p* < .001).

### Age and Partner-Related Neural Responses to Fairness

A main focus of our study was to characterize how fairness-related brain function may differ across age groups and partner similarity. To answer this question, we conducted both whole-brain activation and effective network connectivity analyses. For whole-brain activation analyses, we did not find any brain regions demonstrating a significant interaction between partner similarity and participant age group. However, network connectivity analyses revealed a significant interaction between participant age group and partner similarity (contrast: [young_similar-young_dissimilar] > [old_similar-old dissimilar]) in fairness-related connectivity of the DMN (Figure 3). Specifically, younger adults exhibited stronger fairness-modulated DMN-ACC connectivity when interacting with a similar versus dissimilar partner, whereas older adults demonstrated the opposite pattern (i.e., stronger DMN-ACC connectivity when interacting with a dissimilar versus similar partner; contrast = [young_similar-young_dissimilar] > [old_similar-old dissimilar], peak MNI_xyz_ = −13.3, 34, 27.8; cluster = 26 voxels, p = .023, corrected). We did not find any significant interaction between participant age and partner similarity in ECN network connectivity.

**Figure 3.**
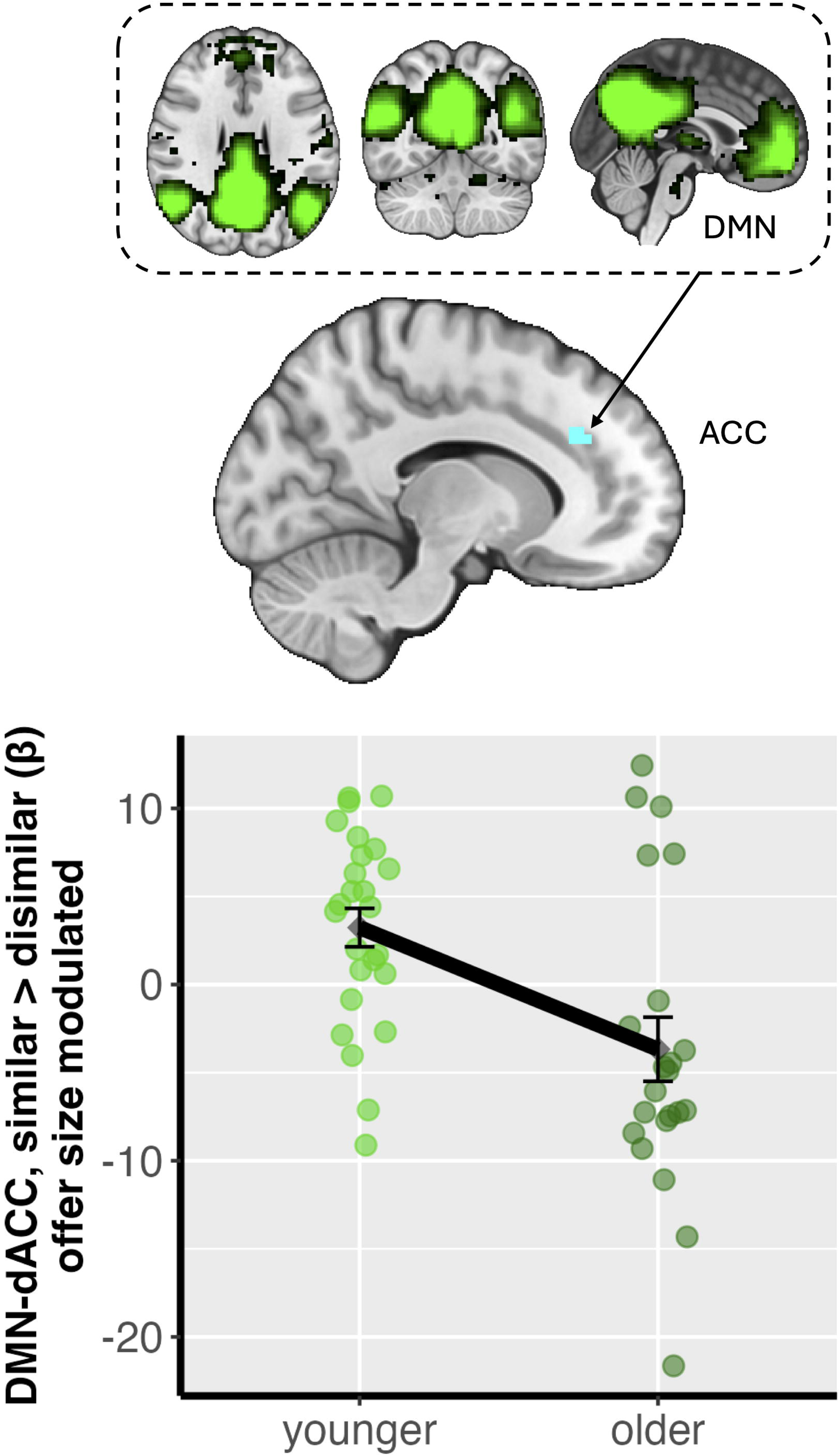
Modulation of Default Mode Network (DMN)-Anterior Cingulate Cortex (ACC) connectivity by age group and partner similarity. The x-axis represents the two Age Groups (25 Younger Adults: 18-35 years; 22 Older Adults: 65-80 years). The y-axis displays the individual-level parameter estimates (beta) reflecting offer size-modulated DMN-ACC connectivity difference between the ‘similar partner’ and ‘dissimilar partner’ conditions (similar > dissimilar, ACC MNIxyz = −13.3, 34, 27.8; cluster = 26 voxels, p = .023). Each dot represents a single participant’s mean connectivity value for this contrast. Error bars indicate the standard error of the mean for each age group. A positive value on the y-axis signifies stronger offer size-modulated DMN-ACC connectivity when interacting with a similar partner compared to a dissimilar partner. Brain image is thresholded and corrected for multiple comparisons using an initial cluster-forming threshold of z > 3.1 followed by a whole-brain corrected cluster-extent threshold of p < 0.05.

In addition to the interaction between age group and partner similarity, we were interested in whether behavior-level individual differences in sensitivity to fairness norm violations further modulated task-based connectivity. To test this, we first estimated each participant’s sensitivity to fairness norm violations using participant-specific slopes from a multilevel logistic regression model (i.e., the sum of group-level fixed offer size effect and a participant’s unique corresponding random slope). A steeper positive slope would indicate a greater sensitivity to the violation of a participant’s fairness norm. We then modeled these participant-specific fairness sensitivity slopes as a covariate into our group-level models for DMN and ECN connectivity. We found a significant effect of fairness norm violation sensitivity on offer size-modulated ECN-mPFC connectivity as a function of age group and partner similarity ([older_similar-older_dissimilar] > [young_similar-young_dissimilar], MNI_xyz_ =4.58, 60.7, 11.7; cluster = 23 voxels, p = .029, corrected) (Figure 4): that younger adults’ behavioral sensitivity to fairness was negatively associated with the difference in ECN-mPFC connectivity when interacting with similar vs. dissimilar partners, whereas older adults showed a positive association. That is, younger adults who were more sensitive to fairness norm violations showed reduced ECN-mPFC connectivity when interacting with similar partners compared to dissimilar partners, whereas older adults demonstrated the reversed relationship, such that those more sensitive to fairness norm violations showed heightened ECN-mPFC connectivity with similar partners relative to dissimilar partners.

**Figure 4.**
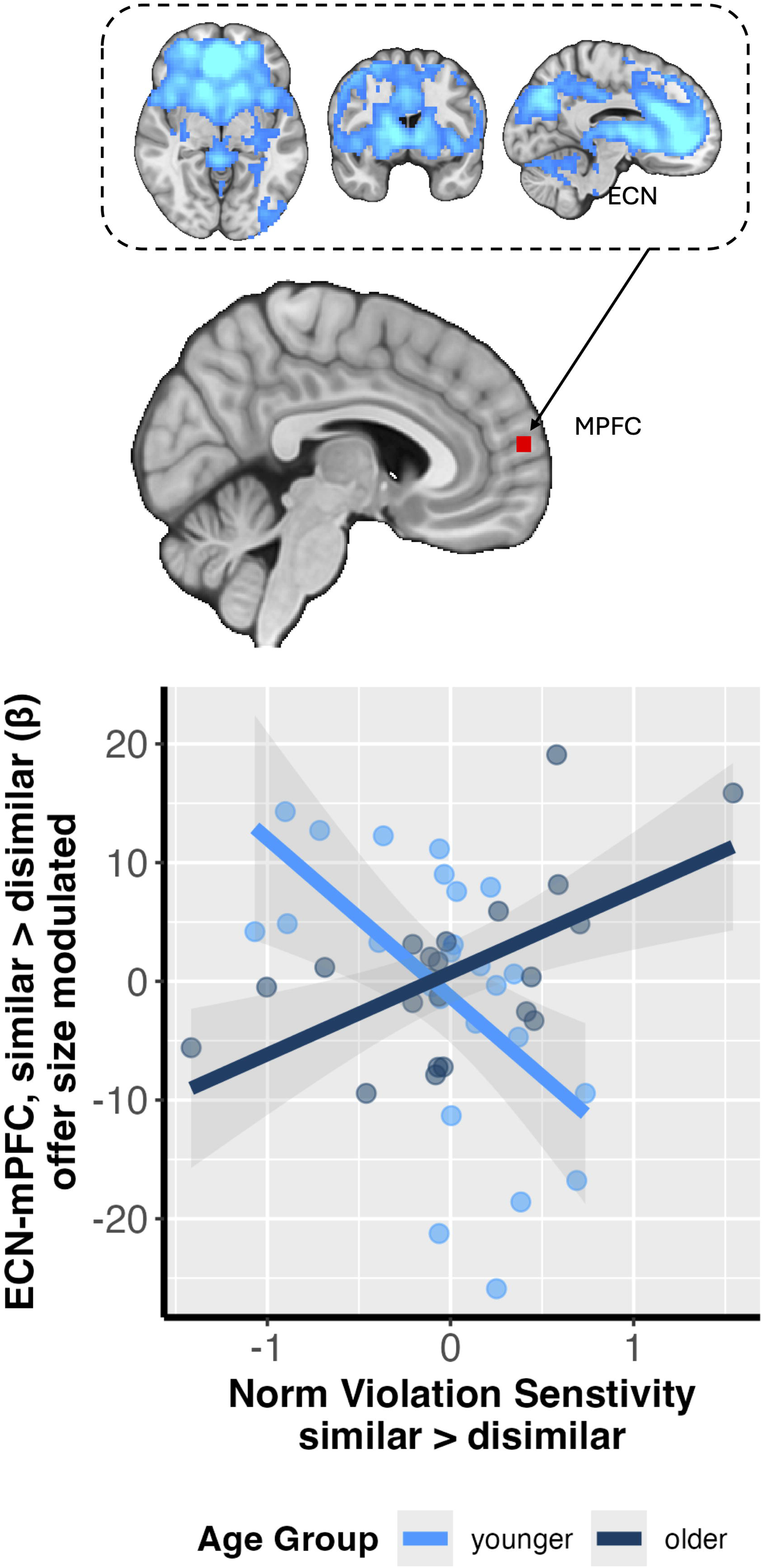
Interaction between sensitivity to fairness norm violations and age on Executive Control Network (ECN)-mPFC connectivity. The x-axis represents individual participant’s norm violation sensitivity difference between similar and dissimilar partners. The y-axis displays the individual-level parameter estimates (beta) reflecting offer size-modulated ECN-mPFC connectivity difference between the ‘similar partner’ and ‘dissimilar partner’ conditions (similar > dissimilar, mPFC MNIxyz =4.58, 60.7, 11.7; cluster = 23 voxels, p = .029). Light blue dots represent individual younger adult participants, while dark blue dots represent individual older adult participants. The solid lines indicate the linear fitted regression lines for each age group. The shaded regions around each fitted line denote the estimated standard error of the fitted line. This figure visualizes the opposing correlations between behavioral fairness sensitivity and neural differentiation in ECN-mPFC connectivity between similar and dissimilar partners across younger and older adults. The brain image is thresholded and corrected for multiple comparisons using an initial cluster-forming threshold of Z > 3.1 followed by a whole-brain corrected cluster-extent threshold of p < 0.05.

## Discussion

The present study investigated how age and social context of partner identity modulate the neurobehavioral correlates of fairness-related economic decision-making. Although younger and older adults did not differ in their choice behavior as they consistently preferred fairer offers regardless of partner similarity, their underlying neural mechanisms diverged. Specifically, younger adults showed increased DMN-anterior cingulate connectivity compared to older adults when considering offers from a similar partner. We also found that context-dependent connectivity between the ECN and mPFC was associated with behavioral sensitivity to violations of fairness norms. ‘Together, these findings indicate that similar choice patterns across age groups were accompanied by different patterns of task-dependent connectivity. These differences may reflect age-related variation in neural processes associated with social decision making, although the present design cannot establish compensatory or adaptive mechanisms.

Our neuroimaging results provide insights into the functional reorganization of brain networks among older adults in relation to social economic decision making. A primary finding was the age-related reversal in DMN-ACC connectivity associated with partner similarity, where younger adults exhibited heightened connectivity when considering offers from a similar partner, while older adults showed the inverse pattern. While intrinsic DMN connectivity has been indicated to often decline with age (Ng et al., 2016), our findings reveal a task-based functional reconfiguration that is critically sensitive to social context. This pattern is consistent with theories like the Default-Executive Coupling Hypothesis of Aging, which posits that older adults may leverage reduced network segregation as an adaptive strategy to support goal-directed cognition (Turner & Spreng, 2015). This neural recalibration was further illuminated by our second key finding: an interaction involving ECN-mPFC connectivity, individual sensitivity to fairness norm violations, age, and the social context of partner similarity. The mPFC is a critical hub for socio-affective processing (Lieberman et al., 2019; Tovar & Chavez, 2021), and its functional community structure is known to become less specialized and more interconnected with other networks during decision-making in older adults (Moussa et al., 2014; Ranjbar-Slamloo et al., 2025). Our results provide evidence for functional consequences for this known age-related reorganization: altered ECN-mPFC connectivity among older adults is meaningfully tied to sensitivity to individual fairness norms, but in a direction opposite to that of younger adults. Taken together, these findings collectively suggest that the mechanisms supporting fairness-related decision-making may be functionally recalibrated as we age in response to social context and individual sensitivity to norm violations. It is possible that older adults for whom such age-related neural changes is less effective or misaligned with the demands of external social contexts and sensitivity to fairness norms may become more vulnerable to financial scams exploiting insensitivity to social cues (e.g., Spreng et al., 2017). However, we note that the current study did not include direct investigation of the association between age-related brain responses to social context and experiences of or risk for fraud. Future research incorporating such assessments of fraud would help to clarify the real-world implications of age and context-dependent neural responses during social decision making.

Notably, the age and social context-dependent patterns of neural activity we found stood in stark contrast to our behavioral findings, which showed no significant differences in choice behavior between age groups or partner identities. This dissociation between behavioral outcomes and their underlying neural substrates is a well-documented characteristic of cognitive aging and is broadly consistent with theories of neural compensation and functional reorganization (Morcom & Johnson, 2015). Previous research has demonstrated that older adults can achieve task performance comparable to or even better than their younger counterparts by recruiting different or additional neural resources (e.g., Bagarinao et al., 2019; Knights et al., 2024). The age-dependent opposing patterns of DMN-ACC and ECN-mPFC connectivity that we observed may represent such a compensatory shift. While younger adults appear to engage these networks in a manner sensitive to partner similarity, older adults may successfully regulate their behavior to align with social context and fairness norms as younger adults do by engaging these same networks in a different manner. This suggests that older adults can preserve behaviors similar to those of younger adults through different patterns of connectivity of the DMN and ECN.

While our findings illustrate a brain-behavior dissociation wherein older adults recruit distinct neural strategies to achieve normative social decision making, several limitations should be noted. First, we acknowledge that our sample size is relatively modest, which may affect the generalizability of our results and the statistical power to detect more subtle effects (Baranger et al., 2023). Second, our cross-sectional design precludes inferences about longitudinal aging trajectories, as the observed age differences could reflect cohort effects rather than the aging process per se (Argiris et al., 2021). Third, while we observed differences based on similarity between participants’ and partners’ age we did not collect subjective assessments of how similar participants felt to the partners. Finally, we note that the observed interaction between participant age, partner similarity, and behavioral sensitivity to fairness may alternatively reflect an interaction between partner age and fairness sensitivity. That is, the neural patterns we attribute to age-related differences in decision making could instead—or additionally—be driven by characteristics of the proposing partner, such as their age. Future work could adjudicate between these interpretations by systematically measuring and modeling other dimensions of perceived similarity between participants and their partners (e.g., social closeness, gender, shared values, or group membership).

## Conclusion

In summary, the present study advances our understanding of how aging shapes the neural mechanisms underlying fairness-related social decision-making. Behaviorally similar fairness decisions in younger and older adults were accompanied by different patterns of DMN-ACC and ECN-mPFC connectivity across partner contexts. These findings identify candidate network-level correlates of age and social context but do not establish compensation or distinguish participant-age from partner-age effects. By integrating behavioral, neural, and individual difference measures, our work lays the foundation for a more nuanced model of socio-cognitive aging that bridges brain function with real-world behavior in social interactions.

## Funding and data availability

This work was supported by funding from the National Institute on Aging (R01-AG067011 to D.V.S, T32AG066598 to K.H., and F31AG085934 to J.B.W.), a College of Liberal Arts Research Award (to D.V.S.), National Institute of Mental Health (R15-MH122927 to D.S.F.). This work was also partially supported by a Pilot Grant from the Scientific Research Network on Decision Neuroscience and Aging (NIH R24-AG054355 to D.V.S). Our statistical analysis plan was not pre-registered. De-identified data is available on OpenNeuro (https://doi.org/10.18112/openneuro.ds003745.v2.0.2) and described in Smith et al., 2023. Code is available at https://github.com/DVS-Lab/srndna_ug_public.

