## Supplementary material for "Older Adults Show Altered Default Mode and Executive Control Network Connectivity during Fairness Decisions": Table 1

**Table 1. Models and Estimates (fixed effects, *p<0.001)**

| **Models**  **(factors included)** | **Model 1**  **(offer, age, similarity)** | **Model 2**  **(offer, age)** | **Model 3**  **(offer, similarity)** | **Model 4**  **(offer, age as control)** | **Model 5**  **(offer, similarity as control)** | **Model 6**  **(offer)** |
| --- | --- | --- | --- | --- | --- | --- |
| **Regressors** | **Beta Estimates**  **(Standard Error)** | | | | | |
| offer * age * similarity | Failed to converge |  |  |  |  |  |
| offer * age |  | -0.144  (0.305) |  |  |  |  |
| offer * similarity |  |  | -0.057  (0.047) |  |  |  |
| offer |  | 1.721*  (0.218) | 1.686*  (0.172) | 1.660*  (0.17) | 1.655*  (0.17) | 1.655*  (0.170) |
| age |  | 1.523  (2.133) |  | 0.969  (1.783) |  |  |
| similarity |  |  | 0.271  (0.239) |  | 0.022  (0.123) |  |
| intercept |  | -5.717*  (1.476) | -5.152*  (1.092) | -5.463*  (1.364) | -5.024*  (1.089) | -5.013*  (1.087) |
| **Observations** |  | 4439.00 | 4439.00 | 4439.00 | 4439.00 | 4439.00 |
| **Log Likelihood** |  | -943.05 | -942.59 | -943.15 | -943.28 | -943.30 |
| **AIC** |  | 1900.11 | 1899.19 | 1898.31 | 1898.57 | 1896.60 |
| **BIC** |  | 1944.89 | 1943.98 | 1936.69 | 1936.96 | 1928.59 |
| *p<0.001  similarity = partner similarity; offer = offer size; age = participant age | | | | | | |
